# Green synthesis of silver nanoparticles from *Artemisia afra* and *Artemisia annua* leaf extracts: Enhanced *in vitro* anthelmintic activity against *Haemonchus contortus*

**DOI:** 10.64898/2026.10.09.757962

**Authors:** Evouna Madeleine Ines Danielle, Bogning Zangueu Calvin, Ntoumba Agnes Antoinette, Tako Djimefo Alex Kevin, Eya’ane Meva Francois

## Abstract

Rising anthelmintic resistance in *Haemonchus contortus* and other gastrointestinal nematodes poses a critical challenge to livestock parasite management, underscoring the urgent need for novel therapeutic agents. This study evaluated the *in vitro* anthelmintic potential of green-synthesized silver nanoparticles (AgNPs) derived from *Artemisia afra* and *Artemisia annua* leaf extracts against adult worms. Aqueous leaf extracts were utilized to biosynthesize AgNPs by optimizing reaction parameters, including extract-to-silver nitrate ratio, concentration, pH, and incubation time. Phytochemical characterization identified alkaloids, coumarins, tannins, polyphenols, and saponins in both species, with *A. afra* demonstrating markedly higher total phenolic (282 mg GAE/g) and flavonoid (3655.56 mg QE/g) contents than *A. annua*. UV-Vis spectroscopy confirmed nanoparticle synthesis, exhibiting characteristic surface plasmon resonance peaks at 435 nm (*A. afra*-AgNPs) and 456 nm (*A. annua*-AgNPs). Anthelmintic efficacy was assessed using the Adult Worm Motility Assay across a concentration gradient (3–48 µg/mL) over 24 hours. Both AgNP formulations displayed dose-dependent inhibition of motility. Notably, *A. afra*-AgNPs exhibited superior potency, achieving lower IC_50_ values at 12 hours compared to *A. annua*-AgNPs and the commercial anthelmintic levamisole. At 48 µg/mL (12 hours), *A. afra*-AgNPs demonstrated significantly greater motility inhibition than *A. annua*-AgNPs, while surpassing levamisole at 12 µg/mL (18 hours). These findings highlight green-synthesized AgNPs from *Artemisia* species, particularly *A. afra*, as potent, eco-friendly nano-anthelmintic candidates, offering a sustainable approach to mitigate drug-resistant nematode infections in livestock.

## 1. Introduction

Gastrointestinal nematodes, particularly the hematophagous parasite *Haemonchus contortus*, present a major burden to small ruminant livestock production. Infections lead to reduced weight gain, diminished milk production, and elevated mortality rates, resulting in severe economic losses especially in tropical and subtropical regions (Upadhyay et al., 2024). While synthetic anthelmintics have traditionally formed the primary strategy for parasite management, widespread drug resistance has compromised their efficacy, emphasizing the necessity for novel, sustainable, and eco-friendly therapeutic alternatives (Mustafa et al., 2024).

Medicinal plants represent a rich source of bioactive secondary metabolites with anthelmintic potential. The genus *Artemisia* is notably recognized for its phytochemical diversity and historical application in traditional antiparasitic medicine (Cruz-Estupiñan et al., 2025). Species such as *Artemisia annua* and *Artemisia afra* synthesize a broad array of bioactive compounds, including artemisinin, flavonoids, phenolic acids, and terpenoids. However, the therapeutic utility of crude plant extracts is frequently constrained by poor bioavailability, low aqueous solubility, and rapid chemical degradation *in vivo* (Sakyi et al., 2025).

Nanotechnology provides an advantageous approach to overcome these biopharmaceutical limitations. Green-synthesized silver nanoparticles (AgNPs) utilize plant phytochemicals as natural reducing and capping agents, combining botanical biological activity with the favorable physicochemical attributes of nanomaterials (Ali et al., 2021; Pereira et al., 2025). Although prior studies have reported anthelmintic activity of AgNPs biosynthesized from *Artemisia cina* and *Artemisia judaica* against *H. contortus*, a systematic comparison evaluating the adulticidal efficacy of AgNPs derived from *A. afra* versus *A. annua* has not been conducted (Guerrero et al., 2025; Mostafa et al., 2024).

It was hypothesized that green-synthesized AgNPs from both *Artemisia* species would exhibit superior *in vitro* anthelmintic activity against adult *H. contortus* compared to crude extracts and the commercial drug levamisole. Accordingly, the primary objective of this study was to synthesize, optimize, and characterize *A. afra*-AgNPs and *A. annua*-AgNPs, and to quantify their time- and concentration-dependent impacts on adult worm motility. A secondary objective was to correlate observed anthelmintic potency with specific phytochemical constituents, particularly total phenolic and flavonoid contents. Developing accessible, plant-derived nano-anthelmintics provides a viable option to combat drug-resistant helminths, addressing a critical need in veterinary parasitology and livestock health management (Castagna et al., 2021).

## 3. Materials and methods

### 3.1 Study design and plant material collection

A six-month study was conducted at the Faculty of Medicine and Pharmaceutical Sciences, University of Douala (Douala, Cameroon). Dried leaves and stems of *Artemisia annua* and *Artemisia afra* were procured from the Artemisia House in Bafia, Cameroon, under batch numbers 24112201 and 24101702. The dried plant material was ground into a fine powder using an electric mill and stored in airtight containers at room temperature 25±2 °C, protected from light.

### 3.2 Preparation of aqueous extracts

Aqueous extracts were prepared using a modified infusion method (Shinyuy et al., 2023): Briefly, 5 g of powdered plant material was steeped in 100 mL of distilled water at 80°C for 5 min. Following cooling to room temperature, the mixture was filtered through Whatman No. 1 filter paper. The filtrate was dried at 40 °C in an oven to obtain a dry powder extract. The extraction yield was determined as the percentage of dry extract mass relative to the initial dry plant mass (Equation 1):

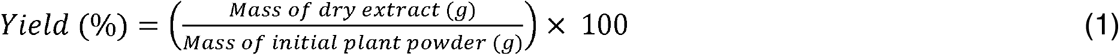

### 3.3 Qualitative and quantitative phytochemical screening

Qualitative phytochemical screening was performed using standard colorimetric and precipitation assays and spectrophotometric methods were employed to quantify total phenolic content (TPC), total flavonoid content (TFC), and total tannin content (TTC) (□icolea et al., 2025). TPC was determined via the Folin-Ciocalteu method using gallic acid as the standard (Atanassova et al., 2011). Extract solutions (0.5 mL, 1 mg/mL) were mixed with 2.5 mL of 10% (v/v) Folin-Ciocalteu reagent and 2.0 mL of 7.5% (w/v) sodium carbonate (Na_2_CO_3_). After a 30-min incubation at room temperature, absorbance was measured at 765 nm using a Shimadzu UV-1800 spectrophotometer. Results are expressed as milligrams of gallic acid equivalents per gram of dry extract (mg GAE/g). TFC was measured using the aluminum chloride (AlCl_3_) colorimetric method with quercetin as the calibration standard (Chang et al., 2002). Extract solution (1.0 mL) was combined with 1.0 mL of 2% AlCl_3_ in methanol, incubated for 10 min, and read at 430 nm. Results are expressed as milligrams of quercetin equivalents per gram of dry extract (mg QE/g). TTC was assessed via the Folin-Ciocalteu assay using tannic acid as the reference standard (Saxena et al., 2013). Results are reported as milligrams of tannic acid equivalents per gram of dry extract (mg TAE/g).

### 3.4 Green synthesis and optimization of silver nanoparticles

Silver nanoparticles (AgNPs) were synthesized by reacting aqueous leaf extracts with silver nitrate AgNO_3_, Sigma-Aldrich, ≥99 % purity). Synthesis parameters were systematically optimized by varying the extract-to-AgNO_3_ volume ratio (1:5, 2:5, 3:5, 4:5, and 5:5), the silver nitrate concentration (0.001 M, 0.01 M, 0.1 M, and 0.5 M), the reaction mixture’s pH (2, 4, 6, 7, 8, and 10, adjusted with 0.1 M HCl or NaOH), and the incubation time at room temperature (5, 10, 15, 20, 30, and 60 minutes) (Abdelghany et al., 2018). Optimal synthesis conditions were established as follows: *A. afra*; 1:5 extract-to-AgNO_3_ ratio with 0.01 M AgNO_3_, at its natural pH of 6.91, and 15 minutes of incubation. *A. annua*; 4:5 ratio, 0.01 M AgNO_3_, pH of 7.4, and 60 minute incubation. Nanoparticle formation was visually indicated by a color shift from pale yellow to dark brown and confirmed by UV-Visible spectrophotometric scanning across 200–1000 nm to record surface plasmon resonance (SPR) peaks (Ijaz Hussain et al., 2011). Formed AgNPs were collected by centrifugation at 12,000 rpm for 20 min, washed three times with distilled water to remove unreacted ions and residual phytochemicals, and resuspended in phosphate-buffered saline (PBS, pH 7.4).

### 3.5 In vitro anthelmintic activity assay

Anthelmintic efficacy was evaluated using the Adult Worm Motility Assay (AWMA) on adult *Haemonchus contortus* (Ferreira et al., 2013). Live adult worms were harvested from the abomasum of naturally infected sheep freshly slaughtered at a local abattoir in Douala. Abomasa were opened along the greater curvature, washed with warm PBS (37°C), and intact, motile male and female worms were transferred to PBS containing 1% glucose at 37°C. Five active adult worms were distributed per well in 24-well plates, with three independent replicates per treatment group. The experimental treatments comprised green-synthesized Aa-AgNPs and An-AgNPs, crude aqueous extracts of A. afra and A. annua, a silver nitrate solution (0.01 M), levamisole as a positive control, and PBS as a negative control. All treatments were evaluated across five concentrations: 3, 6, 12, 24, and 48 µg/mL (dry weight equivalent). Plates were incubated at 37°C under 5% CO2 in a humidified atmosphere for 24 h. Worm motility was recorded at 0, 6, 12, 18, and 24 h post-exposure by observing individual movement under a stereomicroscope for 30 s. Worms were classified as motile (spontaneous movement) or immotile (complete paralysis). Mortality percentage was calculated using Equation 2:

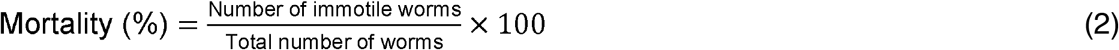

### 3.6 Data analysis and statistical methods

Dose-response curves were constructed by plotting mortality percentages against log_10_ treatment concentrations at each time point. Half-maximal inhibitory concentrations (IC_50_) were evaluated using sigmoidal and Weibull regression models (Ritz et al., 2015). The logarithmic model was excluded from final analysis due to non-convergence and aberrant value fitting on extreme datasets.

Statistical evaluations were conducted using one-way ANOVA (for single-factor comparisons) and two-way ANOVA (for concentration and time interaction effects), followed by Tukey’s and Šídák’s *post-hoc* multiple comparisons tests, respectively (McHugh, 2011). Statistical significance was set at *P* < 0.05. All statistical procedures and curve fittings were executed using GraphPad Prism (version 9.0). Data are presented as mean ± standard deviation (SD) of three independent experimental runs.

## 4 Results

### 4.1 Extraction yields and phytochemical composition

Significant variations in extraction yields were observed between the two species (*p* < 0.05). *Artemisia annua* yielded 34.41%, whereas *Artemisia afra* yielded 7.48% (Table 1). This four-fold difference reflects a higher proportion of water-soluble hydrophilic constituents, such as polysaccharides, in *A. annua* under the aqueous infusion conditions applied (80°C for 5 min) (Shinyuy et al., 2023). Qualitative screening revealed the presence of alkaloids, coumarins, tannins, polyphenols, and saponins in both species (Cruz-Estupiñan et al., 2025). Flavonoids were detected exclusively in *A. afra*, consistent with reports highlighting its elevated levels of luteolin, apigenin, and related glycosides (Septembre-Malaterre et al., 2020). Anthocyanins were absent in both species. Quantitative analysis demonstrated significant disparities in secondary metabolite contents (*p* < 0.05). *A. afra* contained 2.1-fold higher total phenolic content and 5.2-fold higher total flavonoid content compared to *A. annua*. Total tannin content was also higher in *A. afra*.

**Table 1.** Extraction yields and phytochemical composition of *Artemisia afra* and *Artemisia annua* aqueous extracts.

| Parameter | *A. afra* | *A. annua* |
| --- | --- | --- |
| Extraction yield (%) | 7.48 | 34.41 |
| Total phenolic content (mg GAE/g) | 282 ± 12 | 133 ± 9 |
| Total flavonoid content (mg QE/g) | 3656 ± 146 | 700 ± 32 |
| Total tannin content (mg TAE/g) | 45 ± 4 | 22 ± 2 |

Values are presented as mean ± standard deviation of three independent measurements. GAE: gallic acid equivalents; QE: quercetin equivalents; TAE: tannic acid equivalents.

The elevated phenolic, flavonoid, and tannin concentrations in *A. afra* are directly relevant to anthelmintic potential, as these compound classes disrupt parasitic neuromuscular function, energy metabolism, and cuticle integrity (Verma et al., 2023; Olanrewaju et al., 2023). These metabolic distinctions reflect ecological adaptations: *A. afra*, a perennial shrub native to southern Africa, accumulates constitutive phenolics in response to solar radiation and herbivory (Yin et al., 2025), whereas *A. annua*, an annual herb native to temperate Asia, channels primary carbon resources toward artemisinin and sesquiterpenoids (Shen et al., 2018). Observed extraction yields align with reported literature ranges of 25–40% for *A. annua* (Ferreira et al., 2024), and 5–10% for *A. afra* (Shinyuy et al., 2025).

### 4.2 Nanoparticle synthesis and characterization

Synthesis of silver nanoparticles (AgNPs) was visually indicated by a color transition from pale yellow to dark brown, corresponding to surface plasmon resonance (SPR) (Ijaz Hussain et al., 2011). Reaction kinetics differed between species: *A. afra* extracts induced complete color transformation within 5 min, whereas *A. annua* required 15 min, indicating faster reduction kinetics mediated by the higher flavonoid and phenolic content of *A. afra* (Vijayaram et al., 2024). UV-Visible spectrophotometric analysis confirmed AgNP formation. *A. afra*-synthesized AgNPs displayed a distinct SPR peak centered at 435 nm, characteristic of spherical silver nanoparticles (Jain et al., 2007). Absorbance intensity at 435 nm increased from 1.0 at 5 min to 3.40 at 15 min under optimized conditions (1:5 extract-to-AgNO_3_ ratio, 0.01 M AgNO_3_, pH 6.91). plateauing thereafter. Spectral stability observed at 15, 30, and 60 min confirmed nanoparticle stability without significant aggregation.

**Figure 1.**
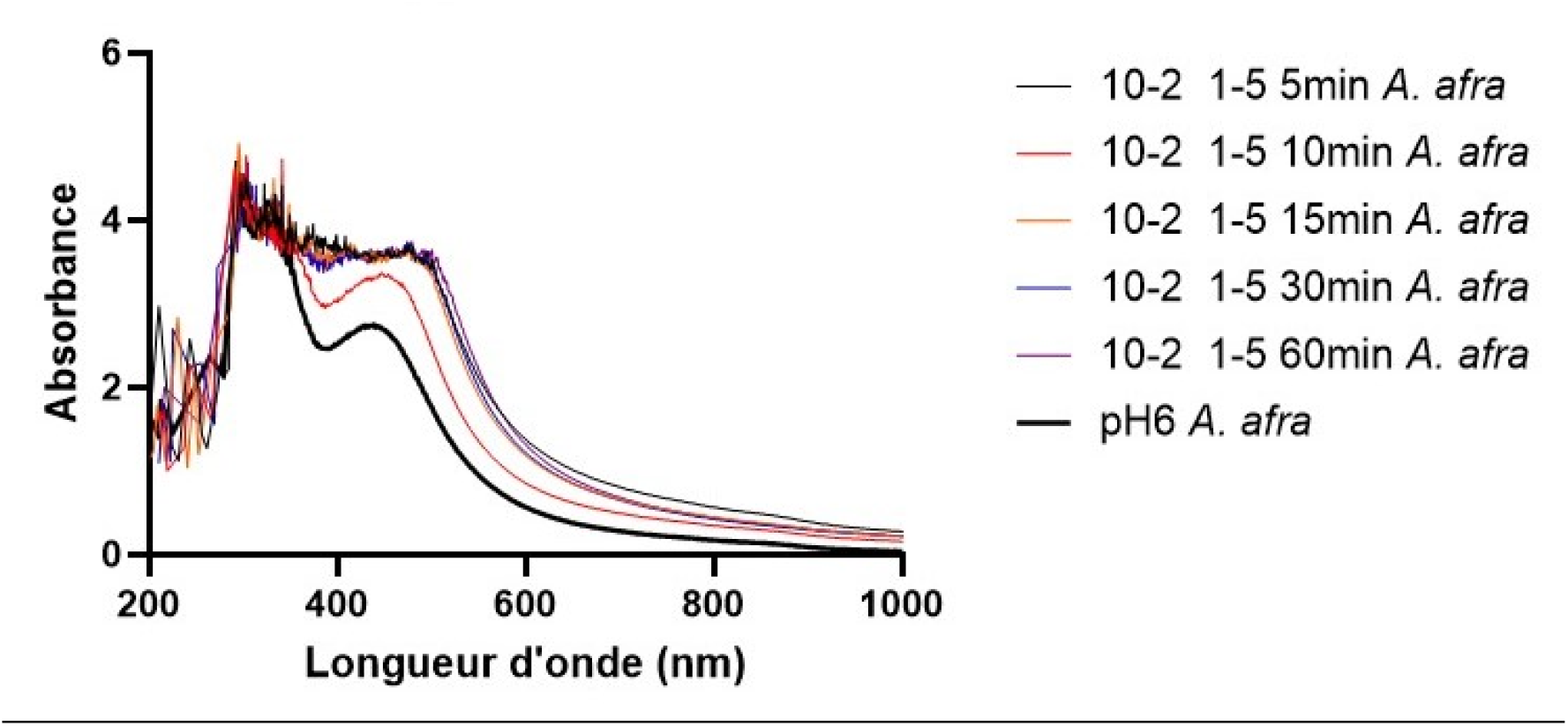
UV-Vis absorption spectra of silver nanoparticles synthesized from *Artemisia afra* extract at different reaction times (5 to 60 minutes) compared to the control extract at pH 6 *A. annua*-synthesized AgNPs (*An*-AgNPs) exhibited an SPR absorption maximum at 456 nm. This red-shift relative to *A. annua*-synthesized AgNPs indicates larger particle dimensions or modified morphological characteristics (Mock et al., 2002). Reaction kinetics for *An*-AgNPs were slower, reaching maximum absorbance (3.69) at 60 min under optimized conditions (4:5 ratio, 0.01 M $\text{AgNO}_3$, pH 7.4). Broader initial SPR bandwidths for *An*-AgNPs suggest a wider size distribution relative to *A, annua*-synthesized AgNPs (Dawadi et al., 2021).

**Figure 2.**
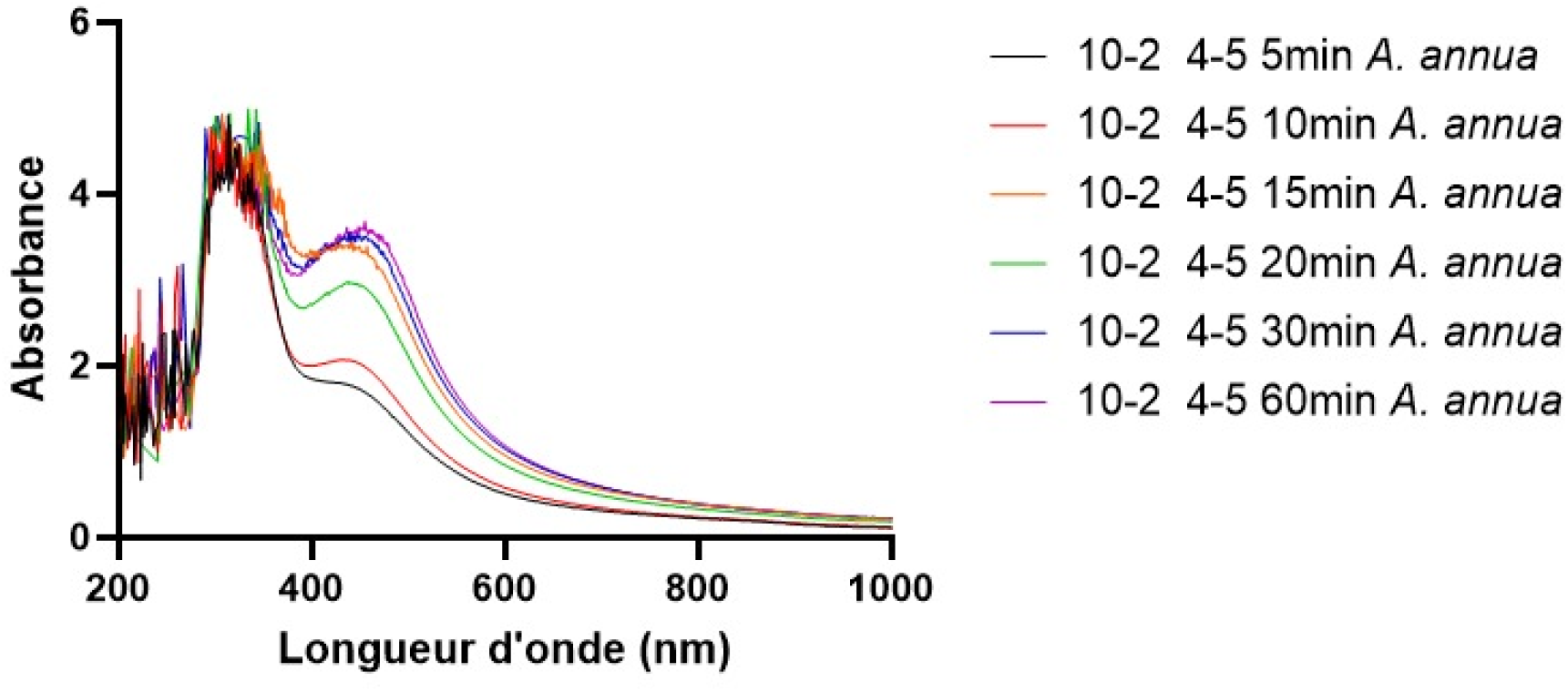
UV-Vis absorption spectra of silver nanoparticles synthesized from *Artemisia annua* extract over time intervals ranging from 5 to 60 minutes Synthesis yields were pH-dependent (Iravani & Zolfaghari, 2013). Optimal synthesis for *A. afra*-synthezised AgNPs occurred at unadjusted pH (6.91), whereas acidic conditions (pH 6.0) reduced yield due to protonation of phenolic hydroxyl groups. For *A. annua*-synthesized AgNPs, maximum synthesis was achieved at pH 7.4, which promotes phenolic deprotonation and enhances electron transfer during Ag^+^ reduction.

**Figure 3.**
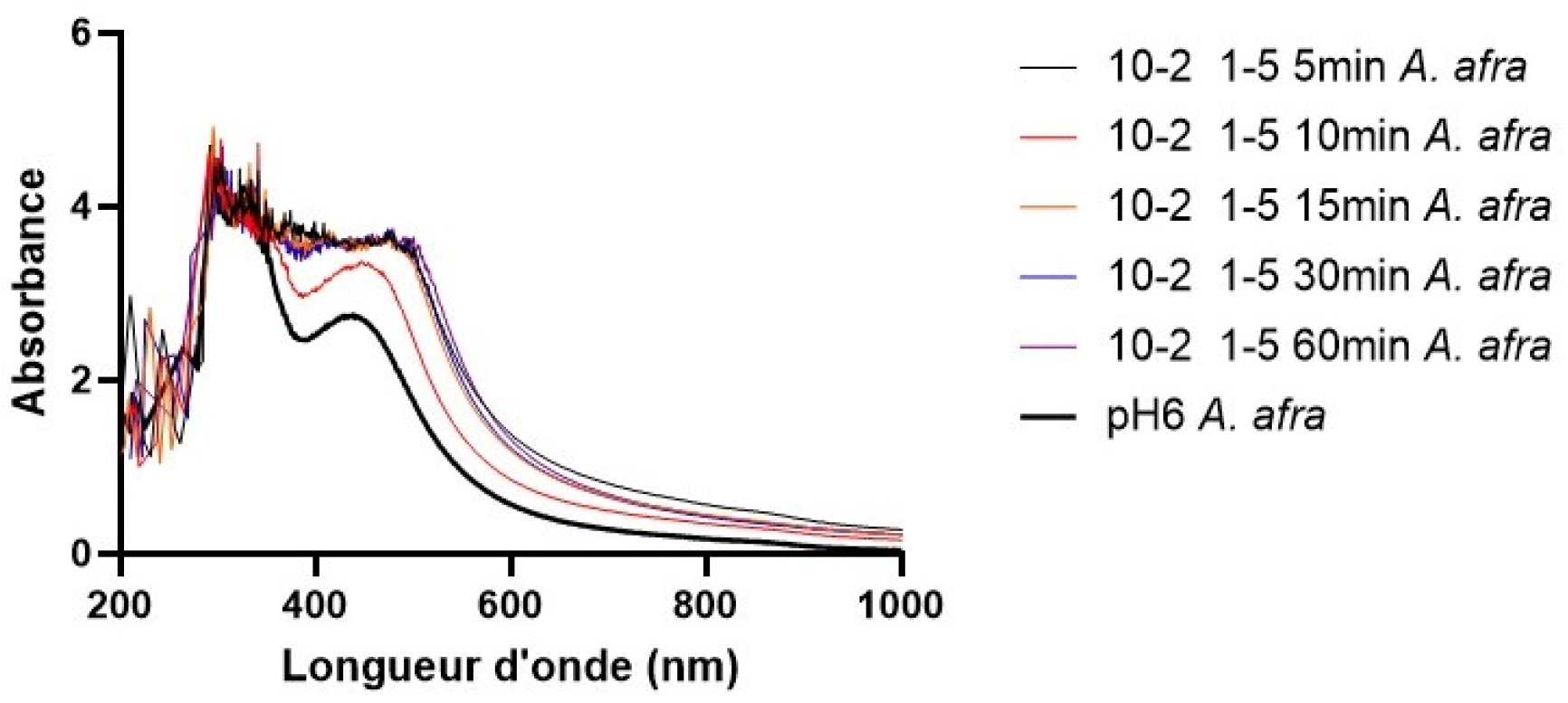
UV-Vis absorption spectra of silver nanoparticles synthesized using *Artemisia afra* extract at different incubation times and pH conditions The rapid reduction rate and defined SPR peaks of *Aa*-AgNPs are attributed to its higher total phenolic and flavonoid content, as flavonoids possess low redox potentials and chelate metal ions effectively during nucleation (Hussain et al., 2019). Slower synthesis in *A. annua* reflects lower reducing phytochemical availability and the presence of non-reducing sesquiterpenoids (Carrillo-López et al., 2016). Absorbance noise observed between 200–350 nm corresponds to unreacted biological macromolecules (Afshinnia et al., 2018). Stable SPR peak intensities over time confirm effective phytochemical capping, preventing nanoparticle aggregation and maintaining physical stability (Kordy et al., 2022; Khan et al., 2014).

## 5. Discussion

These results carry real weight for developing alternative anthelmintic strategies, especially given the mounting resistance crisis in gastrointestinal nematodes. The fact that green-synthesized silver nanoparticles from both Artemisia afra and Artemisia annua showed potent, dose-dependent activity against adult Haemonchus contortus proves that nanotechnology can effectively unlock the bioactive potential of medicinal plants to create agents with genuine practical punch. The standout performance of A. afra AgNPs, with IC_50_ values below those of both A. annua AgNPs and levamisole, makes it clear that a plant’s phytochemical richness directly fuels nanoparticle potency. From a farmer’s perspective in resource-limited settings, these findings offer a tangible, affordable pathway. Using locally grown Artemisia for on-farm nanoparticle production could cut dependence on costly, supply-chain-dependent imported drugs, making parasite control more sustainable and accessible. The rapid synthesis time for A. afra (just 15 minutes) and the mild water-based conditions mean smallholders could adopt this approach with proper training and safety measures in place. Policymakers and veterinary authorities would do well to consider integrating such nanobased tools into integrated parasite management frameworks, particularly where resistance to benzimidazoles and macrocyclic lactones is already widespread (Mederos et al., 2018). several study limitations must be acknowledged. The in vitro AWMA, though a workhorse for initial screening, does not fully mimic the complex gut environment with its pH shifts, enzymes, and microbiota that could alter nanoparticle stability and bioavailability (Zamanian & Chan, 2021). Using adult worms from slaughtered sheep also introduces variability in age, health, and prior drug exposure that might affect baseline motility and susceptibility. (Akhter et al., 2024). Cytotoxicity against mammalian cells was not assessed, a non-negotiable step before any therapeutic use, and relying on just levamisole as a positive control and PBS as a negative might not fully contextualize the activity, as comparisons with ivermectin or albendazole would have strengthened the case.

Looking forward, several logical next steps should be pursued: in vivo trials in sheep or goats to test efficacy, pharmacokinetics, and safety under natural infection, including histopathology of gut tissues and histochemical tracking of nanoparticle distribution; exploring synergy with sub-therapeutic doses of conventional drugs to lower the required nanoparticle load and delay resistance emergence (Zhang et al., 2023); investigating molecular mechanisms through transcriptomic and proteomic work to pinpoint exactly which pathways (neuromuscular, oxidative stress, cuticle synthesis) are disrupted; conducting systematic shelf-life studies under different storage conditions for field readiness; comparative screens across other Artemisia species (e.g., A. absinthium, A. vulgaris) to identify even more potent sources; and scalability investigations to pave the way for commercial production.

## 6. Conclusion

In conclusion, green-synthesized silver nanoparticles derived from *Artemisia afra* and *Artemisia annua* leaf extracts demonstrate potent, dose-dependent anthelmintic activity against adult *Haemonchus contortus*. Notably, *A. afra*-synthesized AgNPs exhibited superior efficacy, outperforming both *A. annua*-AgNPs and the commercial anthelmintic levamisole. This enhanced potency directly correlates with the higher phenolic and flavonoid content of *A. afra*, which serves a dual role by facilitating rapid nanoparticle reduction and enhancing target bioactivity.

